# Evaluation of the antiviral effect of chlorine dioxide (ClO_2_) using a vertebrate model inoculated with avian coronavirus

**DOI:** 10.1101/2020.10.13.336768

**Authors:** Xóchitl Zambrano-Estrada, Carlos Domínguez-Sánchez, Marina Banuet-Martínez, Fabiola Guerrero de la Rosa, Teresa García-Gasca, Luis Prieto-Valiente, Karina Acevedo-Whitehouse

**Author notes:** **Corresponding author:** Karina Acevedo-Whitehouse.

## Abstract

**Background:** The need for safe and effective antiviral treatments is pressing given the number of viral infections that are prevalent in animal and human populations, often causing devastating economic losses and mortality. Informal accounts of anecdotal use of chlorine dioxide (ClO_2_), a well-known disinfectant and antiseptic, in COVID-19 patients has raised concern about potential toxicity, but also raises the question that ClO_2_ might elicit antiviral effects, a possibility that has never been examined *in vivo* in any animal model. Here, we challenged the hypothesis that ClO_2_ decreases the viral load and virus-induced mortality in a vertebrate model. For this, we determined viral load, virus-induced lesions and mortality in 10-day old chick embryos inoculated with 10^4^ mean EID_50_/mL of attenuated Massachusetts and Connecticut avian coronavirus (IBV) strains.

**Results:** The ClO_2_ treatment had a marked impact on IBV infection. Namely, viral titres were 2.4-fold lower and mortality was reduced by half in infected embryos that were treated with ClO_2_. Infection led to developmental abnormalities regardless of treatment. Lesions typical of IBV infections were observed in all inoculated embryos, but severity tended to be significantly lower in ClO_2_-treated embryos. We found no gross or microscopic evidence of toxicity caused by ClO_2_ at the doses used herein.

**Conclusions:** Our study shows that ClO_2_ could be a safe and viable way of treating and mitigating the effects of avian coronavirus infections, and raises the possibility that similar effects could be observed in other organisms.

**Graphical abstract:** 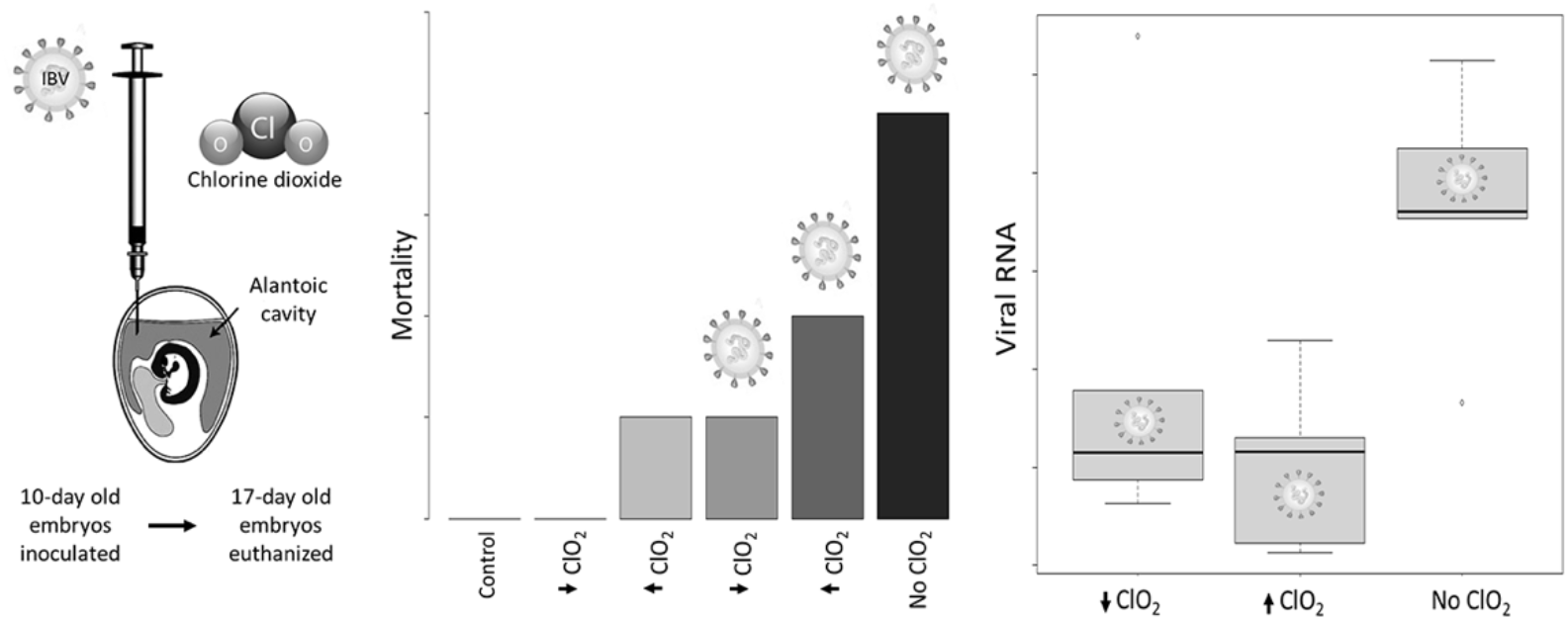

## Background

To date, despite advances in the identification of compounds with antiviral activity, the available repertoire of antiviral drugs is insufficient. Given the number of endemic viruses that impact the health of wild and domestic animals as well as human populations the need for effective and safe antivirals is urgent. The COVID19 pandemic has evidenced this need; since its start in January 2020, a steep rise in cases and associated mortality, mounting pressure on health services worldwide, and a marked damage to the economy of most countries [1, 2] warrants rigorous, unbiased investigation of compounds with potential antiviral action.

The surge in COVID19 cases and the lack of effective treatments have led to a growing number of informal reports and testimonials across social networks regarding the oral and intravenous use of chlorine dioxide (ClO_2_) solution as an efficacious treatment of COVID19. In an unprecedented move, the Senate of Bolivia authorized the extraordinary use of ClO_2_ to “manufacture, sell, supply and use ClO_2_ to prevent and treat COVID19” (law PL N° 219/2019-2020 CS approved 15 July 2020). This situation has sparked concern from health agencies of various countries regarding the potential toxicity of using this disinfectant as a treatment, given that ClO_2_ has strong oxidative properties. Chlorine dioxide has been known since 1946 for its broad antimicrobial properties [3], and it is commonly used to purify water for human consumption, decontaminate produce and other food items [4], and disinfect surgical instruments [5]. According to the Food and Drug Administration (FDA) of the US, consuming ClO_2_ can lead to adverse effects such as ‘vomiting, diarrhoea, dehydration, abdominal pain, metahemaglobinemia and systemic failures that could potentially lead to death’. However, an impartial review of the scientific peer-reviewed literature reveals few case reports of human patients presenting adverse effects after consumption of ClO_2_, none of which were fatal, and reversed completely after treatment [6–9]. Published studies on ClO_2_ toxicity have reported null to mild effects in adult humans and other animals [10–12] and have stressed that toxicity is strongly dose-dependent [12–14].

There is evidence of virucidal activity of ClO_2_ against echovirus [15], enterovirus [16], poliovirus [17], rotavirus [18], norovirus [19, 20], calicivirus [19], and coronavirus [21]. However, there are few publications that explore antiviral effects *in vitro* and *in vivo*. One of the few published studies found that mice exposed to influenza A virus in an environment that contained aerosolized ClO_2_ had significantly lower mortality than mice that were solely exposed to Influenza A virus [22]. The authors reported that the antiviral effect was due to denaturation of hemagglutinin and neuraminidase glycoproteins, a finding that concurs with previous studies that explain the virucidal mechanism of action of ClO_2_ due to oxidation of amino acid residues that are key for cell entry [23–25]. More recently, the antiviral mechanism of action of ClO_2_ was investigated *in vitro* using pig alveolar macrophages and MARC-145 cells exposed to the porcine reproductive and respiratory syndrome virus (PRRSV1), finding that the chemical also inhibits the synthesis of pro-inflammatory molecules that contribute to the pathogenesis of this disease [26]. The authors concluded that viral synthesis of RNA and proteins was impeded by ClO_2_, leading to a reduction in viral replication. Despite these studies, to date there has been no published report of an *in vivo* assessment of the antiviral effect of ClO_2_. Here, we have investigated the effect of a ClO_2_ solution in 10-day old chick embryos inoculated with attenuated avian infectious bronchitis coronavirus (IBV) Massachusetts and Connecticut vaccine strains, which are pathogenic for birds during their embryonic stages of development [27].

## Results

Avian coronavirus RNA was detected in all of the embryos that were inoculated with the vaccine (n=15, in all cases Cq <40). The viral load decreased significantly with increasing ClO_2_ (p=**0.009** for linear trend; Fig. 1), doses being 2.4 times higher in the untreated embryos. The average viral load of ClO_2_-treated chicks was 10^4.3^/mL, range: 10^3.66^ – 10^5.03^ and of untreated chicks was 10^4.83^/mL, range: 10^4.52^ – 10^5.01^, respectively (Tukey HSD, Group E *vs*. F, p = 0.03). There were no differences in the viral load between both ClO_2_-treated groups (p>0.05).

**Figure 1.**
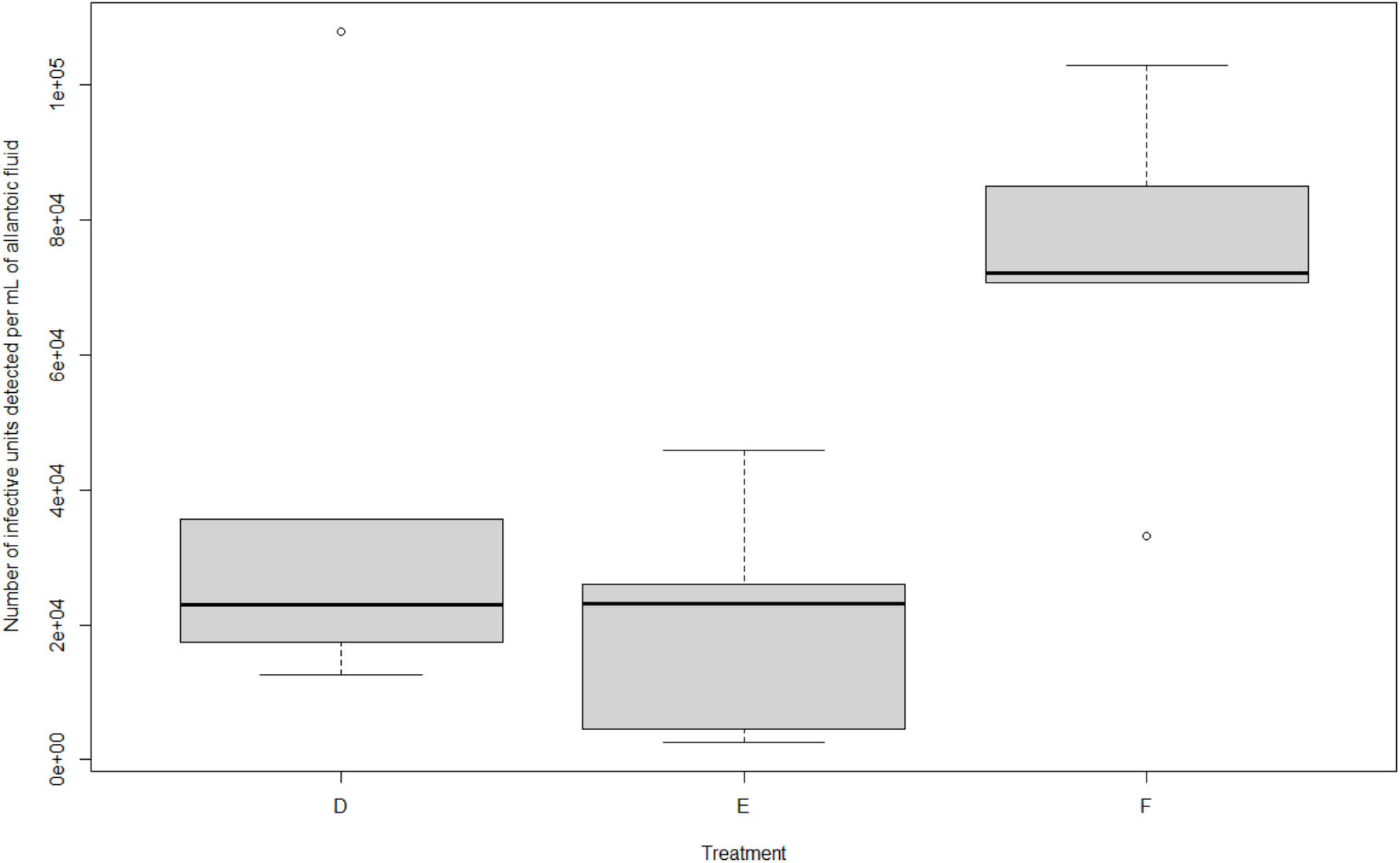
Virus RNA copy number in the allantoic fluid of the infected embryos per treatment group. Virus copy number was calculated by the geometric mean of the triplicate Cq value referred to the standard curve (see methods). Experimental groups D to F contained virus-inoculated embryos (D: Low dose of ClO_2_, E: High dose of ClO_2_, F. Viral control). For details on treatments, see Table 1.

**Table 1.** Experimental groups used to assess the *in vivo* antiviral effect of high and low concentrations of ClO_2_ solution in chick embryos. The table shows the number of embryos included in each group. (LD: Low dose, HD: High dose)

| Group | N | Treatment |
| --- | --- | --- |
| Group A (Experimental control) | 5 | 200 µl of sterile 0.9% saline solution |
| Group B (ClO<sub>2</sub> LD) | 5 | 100 µl of sterile ClO<sub>2</sub> solution (30 ppm<sup>1</sup>) and 100 µl of sterile 0.9% saline solution |
| Group C (ClO<sub>2</sub> HD) | 5 | 100 µl of sterile ClO<sub>2</sub> solution (300 ppm<sup>2</sup>) and 100 µl of sterile 0.9% saline solution |
| Group D (Virus + ClO<sub>2</sub> LD) | 5 | 100 µl of resuspended avian coronavirus vaccine and 100 µl of sterile ClO<sub>2</sub> solution (30 ppm) |
| Group E (Virus + ClO<sub>2</sub> HD) | 5 | 100 µl of resuspended avian coronavirus vaccine and 100 µl of sterile ClO<sub>2</sub> solution (300 ppm) |
| Group F (Viral control) | 5 | 100 µl of resuspended live attenuated avian coronavirus vaccine (Bron Blen® Merial, containing 10<sup>4</sup> of mean embryo infective dose (EID<sub>50</sub>)/mL of coronavirus strains Massachusetts and Connecticut) and 100 µl of sterile 0.9% saline solution |

<sup>1</sup> This concentration is 10 times below the no-observed-adverse-effect level (NOAEL; 3.5 mg/kg per day) determined for ClO<sub>2</sub> (Bercz et al. 1982), considering an embryonic mass of 10 g at the time of inoculation.
<sup>2</sup> This concentration is equal to the NOAEL considering an embryonic mass of 10 g at the time of inoculation.

Mortality differed significantly between virus-inoculated (46.7%) and virus-free embryos (6.7%) (Fisher’s exact p = 0.018), (Fig. 2). In the viral-inoculated groups, 3/10 (30 %) embryos treated with ClO_2_ and 4/5 (80 %) untreated embryos died (Groups D, E and F, respectively). (Fisher’s exact p = 0.119).

**Figure 2.**
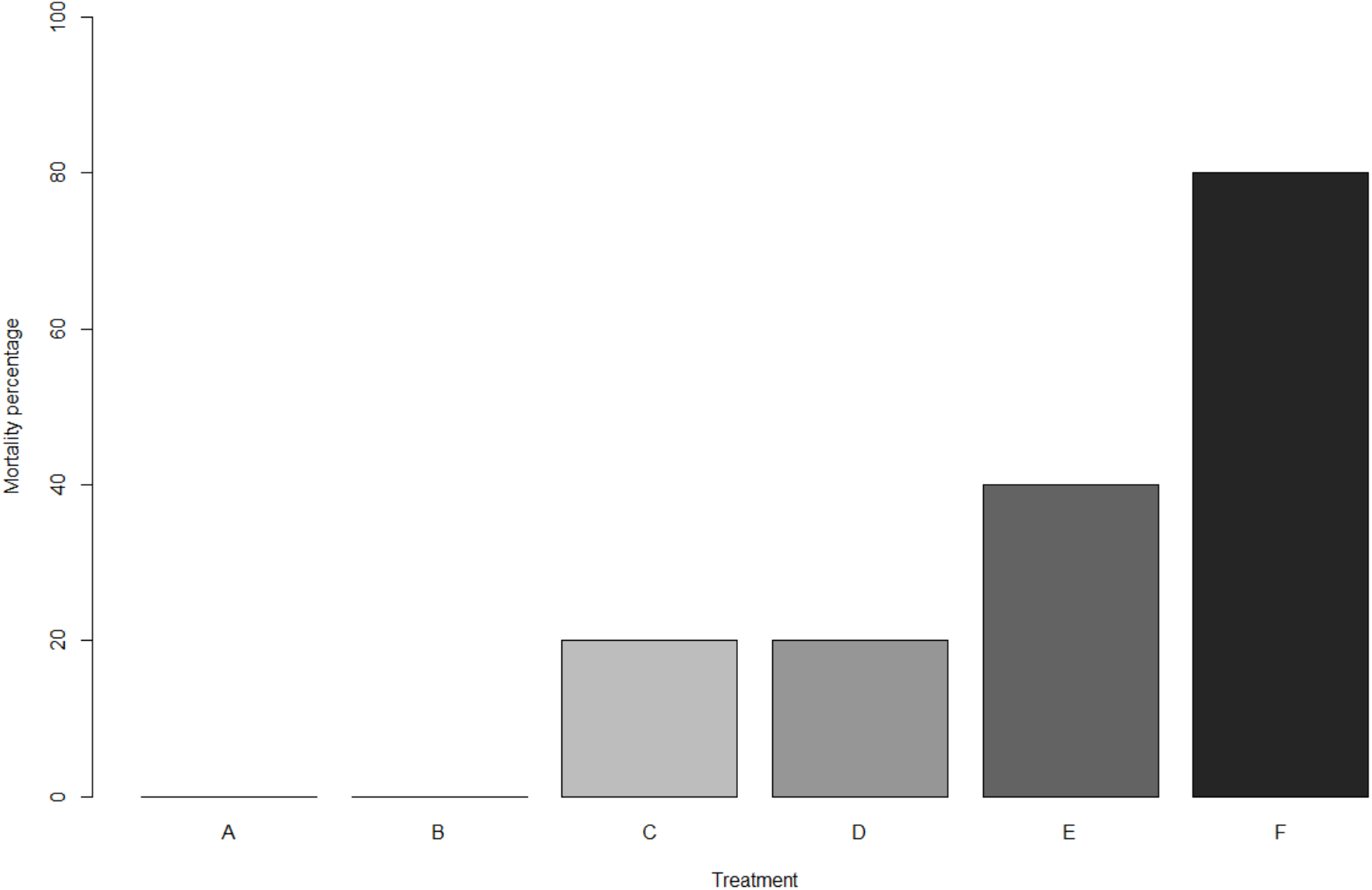
Chick embryo mortality during the experiment. Experimental groups A to C contained virus-free embryos (A: Experimental control, B: Low dose of ClO_2_, C: High dose of ClO_2_); experimental groups D to F contained virus-inoculated embryos (D: Low dose of ClO_2_, E: High dose of ClO_2_, F. Viral control). For details on treatments, see Table 1.

Body mass, embryonic axis length and femur length differed between the virus-inoculated and virus free-groups, regardless of ClO_2_ treatment. All of the virus-inoculated embryos exhibited dwarfing and had, on average, 38% lower mass (t = 21.15, df = 29.40, p = 2.2 × 10^−16^; Fig. 3A), 10% shorter axis length (t = 58.43, df = 29.26, p = 2.2 × 10^−16^; Fig. 3B), and 20% shorter femur length (t = 8.49, df = 23.34, p = 1.4 × 10^−8^; Fig 3C) than the virus-free groups. When analysing growth in the virus-inoculated groups, mass was significantly higher in embryos that were treated with ClO_2_ (t = −2.74, df = 12.98, p = 0.017). Body length of virus-inoculated chicks did not vary according to ClO_2_ treatment (p > 0.1). See supplementary material for photographs of the embryos.

**Figure 3.**
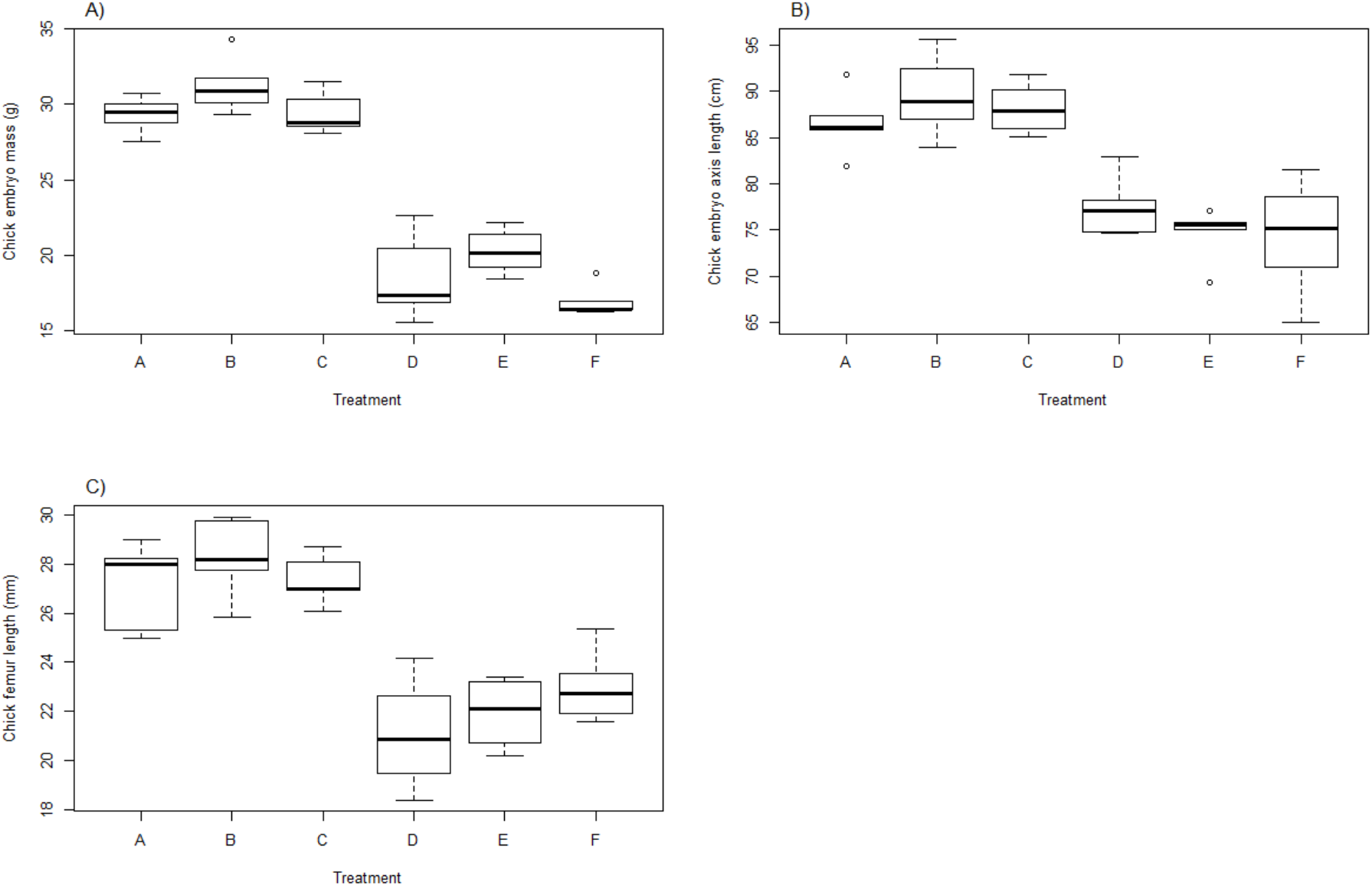
Chick embryo development at the end of the experiment (day 17 of incubation). A) mass, B) body axis, C) femur length. Experimental groups A to C contained virus-free embryos (A: Experimental control, B: Low dose of ClO_2_, C: High dose of ClO_2_); experimental groups D to F contained virus-inoculated embryos (D: Low dose of ClO_2_, E: High dose of ClO_2_, F. Viral control). For details on treatments, see Table 1.

Lesions previously described in embryos infected with avian coronavirus were observed at post mortem examination. Namely, curling, the presence of white caseous material (urates), thickened amnion and allantoic membranes that adhered to the embryos, oedematous serous membranes, epidermal congestion, and subcutaneous haemorrhage (Table 2). Virus-inoculated embryos treated with ClO_2_ had a lower risk of epidermal congestion (RR = 0.4; Wald 95% CI: 0.187-0.855; p = 0.04), haemorrhage (RR = 0.1; Wald 95% CI: 0.016 - 0.642; p = 0.002), curling (RR = 0.019; Wald 95% CI: 0.125 - 0.844; p = 0.017) and thickened membranes (RR = 0; p = 0.003) than untreated infected embryos. All the embryos had a pale liver and mildly congested lungs, regardless of their experimental group. Pale enlarged kidneys were observed in the virus-inoculated groups but not in the virus-free groups, regardless of ClO_2_-treatment (see Table 2).

**Table 2.**
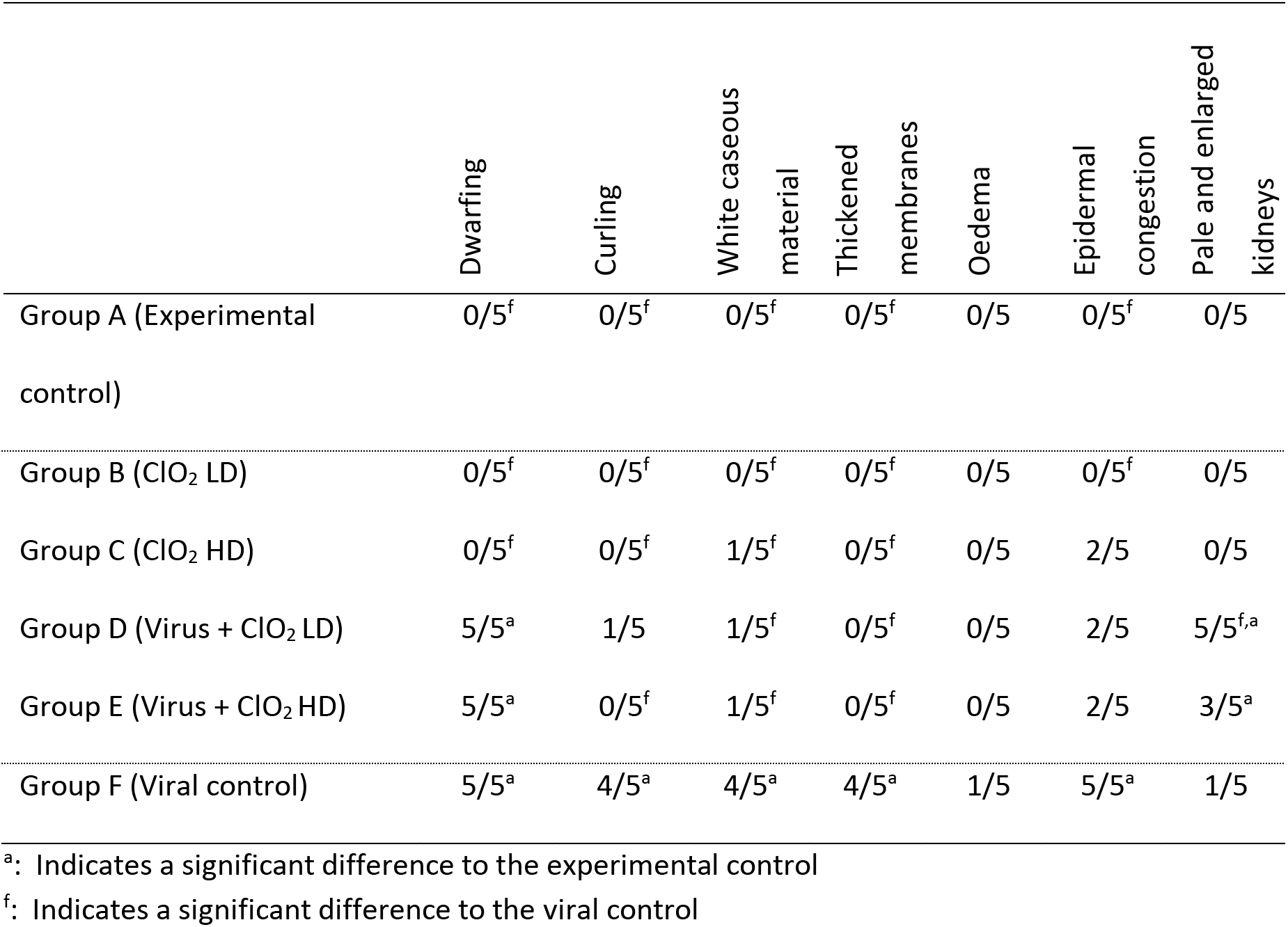
Macroscopic abnormalities observed at necropsy in the chick embryos. The table shows the number of embryos in each group that presented each lesion. (LD: Low dose, HD: High dose)

|  | Dwarfing | Curling | White caseous material | Thickened membranes | Oedema | Epidermal congestion | Pale and enlarged kidneys |
| --- | --- | --- | --- | --- | --- | --- | --- |
| Group A (Experimental control) | 0/5<sup>f</sup> | 0/5<sup>f</sup> | 0/5<sup>f</sup> | 0/5<sup>f</sup> | 0/5 | 0/5<sup>f</sup> | 0/5 |
| Group B (ClO<sub>2</sub> LD) | 0/5<sup>f</sup> | 0/5<sup>f</sup> | 0/5<sup>f</sup> | 0/5<sup>f</sup> | 0/5 | 0/5<sup>f</sup> | 0/5 |
| Group C (ClO<sub>2</sub> HD) | 0/5<sup>f</sup> | 0/5<sup>f</sup> | 1/5<sup>f</sup> | 0/5<sup>f</sup> | 0/5 | 2/5 | 0/5 |
| Group D (Virus + ClO<sub>2</sub> LD) | 5/5<sup>a</sup> | 1/5 | 1/5<sup>f</sup> | 0/5<sup>f</sup> | 0/5 | 2/5 | 5/5<sup>f,a</sup> |
| Group E (Virus + ClO<sub>2</sub> HD) | 5/5<sup>a</sup> | 0/5<sup>f</sup> | 1/5<sup>f</sup> | 0/5<sup>f</sup> | 0/5 | 2/5 | 3/5<sup>a</sup> |
| Group F (Viral control) | 5/5<sup>a</sup> | 4/5<sup>a</sup> | 4/5<sup>a</sup> | 4/5<sup>a</sup> | 1/5 | 5/5<sup>a</sup> | 1/5 |

<sup>a</sup>: Indicates a significant difference to the experimental control
<sup>f</sup>: Indicates a significant difference to the viral control

Microscopic lesions compatible with avian IBV infection were observed in various organs in all virus-inoculated groups (Fig. 4). The severity of the lesions was either similar or slightly lower in the embryos that had been treated with ClO_2_ than in the embryos that did not receive any treatment (Table 3). Two exceptions were the kidneys and the duodenum. In the kidneys, swelling and degeneration of renal tubular epithelium was more common and more severe in the infected chicks that were administered ClO_2_ than in the infected embryos that did not receive ClO_2_, although the former presented mitotic cells. The duodenal villi of the embryos in the IBV-infected groups were longer (ANOVA; F_5,18_ = 5.62, p = 0.003), and their base was wider (ANOVA; F_5,18_ = 13.65, p = 1.39 × 10^−05^) than embryos from the none-infected groups, and they were moderately congested. Duodenal villous atrophy varied amongst groups (ANOVA; F_5,18_ = 5.71, p = 0.003) and *post-hoc* comparisons revealed that the significant differences were E *vs.* B (p = 0.021) and E *vs.* C (p = 0.001). The percent of bursal lymphoid tissue decreased markedly in the virus-inoculated embryos (ANOVA; F_5,12_ = 3.58, p = 0.033; see Fig. 4). Virus-inoculated embryos showed mild apoptosis in the thymus and heterophilic infiltration. Amongst the six experimental groups, all of the embryos examined presented subacute heterophilic bursitis, and pulmonary interstitial multifocal heterophilic foci with congestion (Table 3).

**Figure 4.**
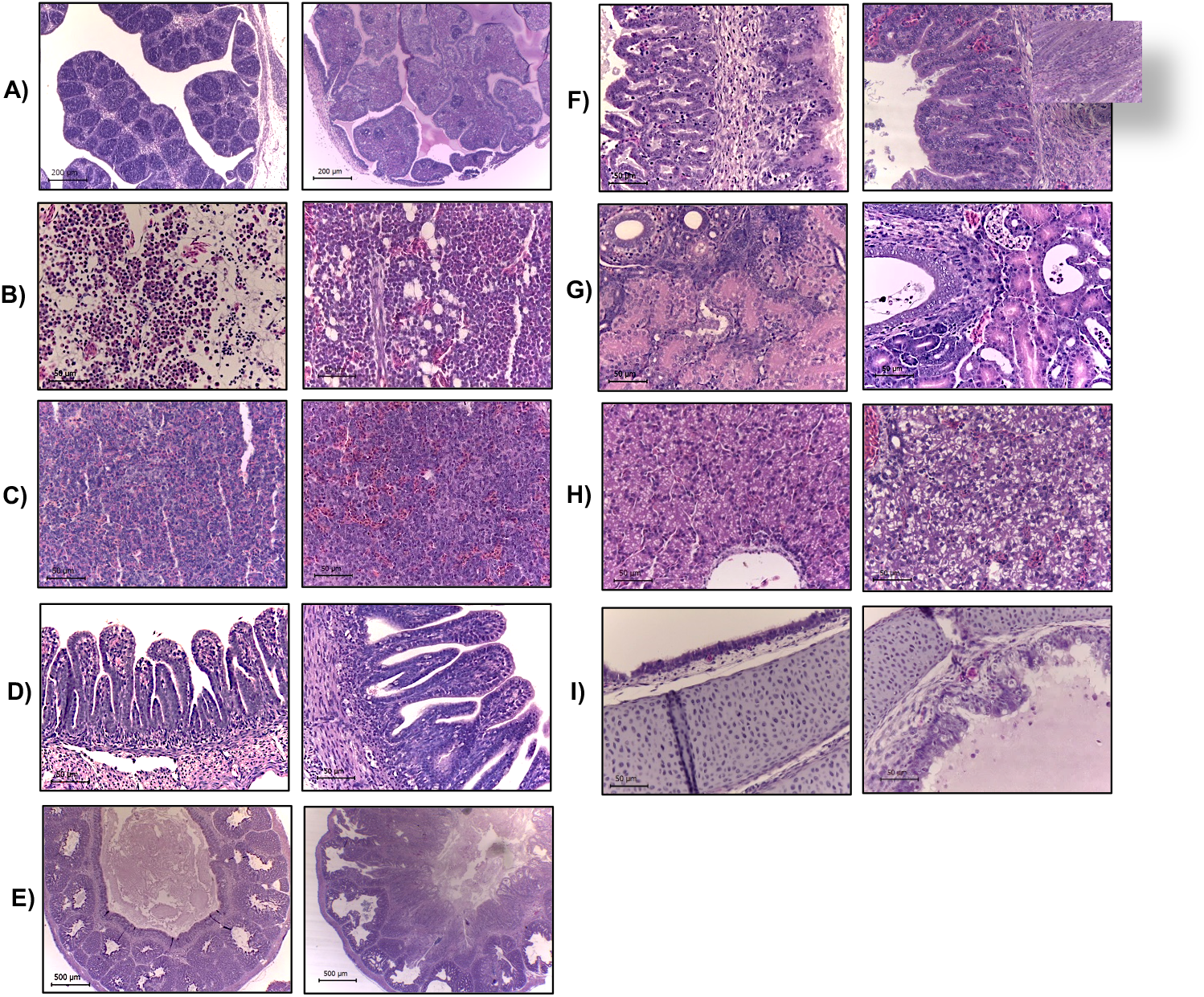
Histology of selected tissues from 17-day-old chick embryos. In each row, microphotograph pairs show H&E-stained representative tissues of non-infected (left) and infected embryos (right), A) Bursa of Fabricio (100X). Infected embryos showed severe lymphoid depletion, with approximately 10% of active lymphoid tissue and abundant heterophils. B) Bone marrow (400 X). Infected embryos showed an increase in erythroid cellularity ne, C) Spleen (400 X), Infected embryos showed reticuloendothelial hyperplasia, D) Duodenum (400 X). Infected embryos showed mild villous atrophy, E) Proventricle (40X). Infected embryos showed epithelial hyperplasia with desquamation and hyaline material, F) Proventricle (400X). Image shows the increased length of the proventricular folds and mitotic cells indicative of epithelial regeneration, G) Kidney (400X). Infected embryos showed swelling, cell detritus and protein within the tubules, and basophilic material compatible with urate crystals, H) Liver (400X). Infected embryos showed glucogenic degeneration, I) Trachea (400X). Infected embryos showed hyperplasia and epithelial degeneration. Scale = 50 μm in all panels except for E (scale = 500 μm).

**Table 3.**
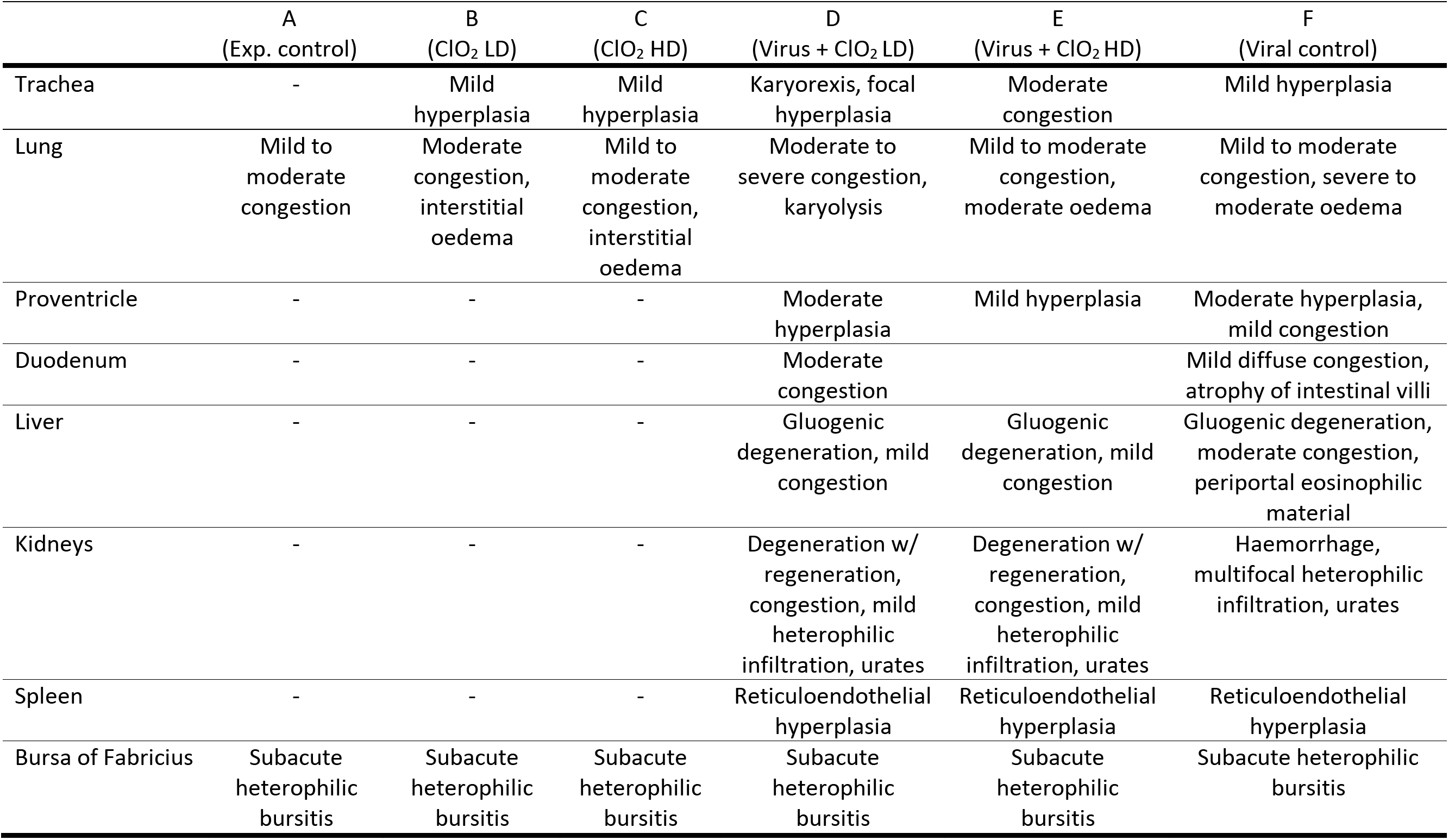
Microscopic abnormalities related to avian coronavirus infection in the chick embryos. (LD: Low dose, HD: High dose)

There were no observable alterations to the architecture and integrity of the tissues of non-infected embryos that were administered ClO_2_ (Experimental groups B and C), nor was any difference in the area (μm^2^) of the thyroid (p>0.1). ClO_2_-treated groups showed a slightly higher myeloid to erythroid ratio in the bone marrow than the experimental control group and the viral control group, although the difference was not statistically significant (ANOVA; F_5,15_ = 2.33, p = 0.094).

## Discussion

We have investigated the antiviral effect of chlorine dioxide (ClO_2_) using avian infectious bronchitis coronavirus (IBV)-infected chick embryos as models. At the concentrations used (30 ppm and 300 ppm), both below the reported NOAEL [13], we observed a reduction in viral titre in infected embryos that were treated with ClO_2_. Furthermore, mortality decreased substantially in the ClO_2_-treated embryos, although alterations to chick development were prevalent regardless of treatment.

Virulent avian IBV strains typically have a burst size of 10 to 100 infective units per cell [28], which can reach peak virus titres of 10^6.5^–10^8.5^ TCID_50_ after 36 hours [29]. The vaccine strains used here have lower replication efficiencies as they are attenuated [30], but they are capable of replicating and causing damage in chick embryos [27]. With 2,000 infective units (200 μl of 10^4^ infective units/mL) inoculated into each embryo, the ClO_2_ treatments decreased viral load 2.4-fold compared to the infected non-treated embryos, representing an average difference of 42,711 infective units. This result could be explained by two non-mutually exclusive mechanisms. Firstly, direct destruction or neutralization of the virions exposed to ClO_2_ could have occurred by denaturing of envelope glycoproteins following oxidation of amino acid residues [22, 23, 25]. Secondly, viral replication efficiency could have decreased due to ClO_2_–induced biochemical changes in the extra- or intracellular milieu, impeding the synthesis of viral RNA and proteins [26, 31].

The effects of reduced viral titres in the infected embryos were evident, with a 50 to 75% reduction in mortality in the treated embryos treated with 300 ppm and 30 ppm of ClO_2_, respectively. However, developmental abnormalities were observed in the majority of the infected embryos, including those that received ClO_2_ treatment. Namely, dwarfing, assessed by body mass, axis length, and femur length were significantly lower in all infected embryos, as expected to occur in IBV infections [32]. In contrast, curling [33, 34] was virtually absent in the embryos that received ClO_2_ after infection. Lesions associated con IBV infection [35, 36] were observed in most of the infected embryos, including those that were administered ClO_2_. However, for most of these lesions, severity was lower or similar in the ClO_2_-treated groups. One exception was in the kidneys, where nephrosis was more severe and frequent in the IBV-infected embryos treated with ClO_2_. Nephrosis is unlikely to have occurred due to ClO_2_ treatment *per se*, given that none of the uninfected embryos that were administered ClO_2_ showed kidney abnormalities. Similarly, atrophy of the duodenal villi was highest in the inoculated group that was administered a high dose of ClO_2_, but absent in the non-infected groups that only received ClO_2_. If duodenal atrophy and nephropathogenicity in IBV-infected embryos are mitigated by inflammatory responses [37], it is possible that the observed damage is due to ClO_2_-driven downregulation of acute inflammation that allowed virion replication in the tubular epithelium and in the duodenal villi. This scenario could be plausible if we consider that ClO_2_-treated pig alveolar macrophages and African green monkey kidney cells infected with the porcine reproductive and respiratory syndrome virus (PRRSV1) downregulate pro-inflammatory cytokines IL-1, IL-6 and TNF-α [26]. There are two drugs, based on chlorite, whose describe mechanism of action might be similar to that elicited by ClO_2_. One is the drug NP001 (a pH-adjusted stabilized form of NaClO_2_) exerts strong anti-inflammatory responses by inhibiting macrophage activation in humans, even after a single dose [38]. In turn, WF10 (tetrachlorodecaoxyde, Cl_4_H_2_O_11_), an oxidizing agent that contains 63.0 mmol/L chlorite and 56.4 mmol/L chloride [39] induces apoptosis of inflammatory cells and downregulates pro-inflammatory genes [40, 41]. More studies will be needed to establish whether the effect of ClO_2_ on IBV-infected chicks is due to similar mechanisms. Interestingly, the affected renal tubules of the chick embryos had mitotic cells, suggestive of regeneration as a reparative response to damage of the renal tubular epithelium [42–44]. This possibility will need to be explored further in the animal model used in our study. Unfortunately, knowledge of IBV pathogenesis, immune responses and tissue reparation in the embryonated egg is limited. In hatched birds, IBV can impact lymphocyte populations by inducing apoptosis, thus impeding virus clearance [45]. We found some evidence of this effect, as the virus-inoculated embryos had a reduced percent of bursal lymphoid tissue.

It is plausible that even though ClO_2_ limited viral replication, given that the embryos were only administered a single dose of ClO_2_ solution, not all virions were eliminated. The viruses that remained viable after administration of the ClO_2_ were able to replicate, yielding lower viral titres, and the damage that they caused to the embryos was less severe and ultimately led to less mortality than in the untreated infected embryos. Repeated administrations of ClO_2_ solution might have reduced the viral load further, plausibly leading to even less virus-induced damage, a possibility that we could not explore in this model. However, an important result of our study was the lack of evidence of tissue damage caused by ClO_2_ itself. Only one (20%) of the uninfected embryos treated with the high dose of ClO_2_ died, and this difference was not statistically significant to the experimental control. It is possible that the death was due to other causes, as a 20% embryo mortality rate is considered normal in aviculture [46, 47].

## Conclusions

Chlorine dioxide had an unequivocal antiviral effect in the model system used in this study, with a reduction in viral RNA, virus-induced lesions and mortality and no evidence of toxicity at the doses used herein. Further studies should aim to test the antiviral effect of ClO_2_ in hatched chicks, which have a more mature immune system, and where repeated administrations are easier to procure. However, it is promising that we found an evident effect against IBV without any significant adverse effect or evidence of toxicity to the chick embryos. Much research needs to be done before it is possible to generalize and extrapolate our findings to other viruses and animal hosts.

## Methods

We aimed to examine whether chlorine dioxide elicits a measurable antiviral effect *in vivo*, and investigate toxicity associated with the administration of this chemical compound. For this, we selected the chicken embryo as a vertebrate model and avian infectious bronchitis coronavirus (IBV) as a virus model. The protocol that is described below was registered at the School of Natural Sciences of the Autonomous University of Queretaro (Mexico). The study was carried out in compliance with the American Veterinary Medical Association (AVMA) guide for humane treatment of chick embryos and approved by the Autonomous University of Queretaro Research Ethics Committee

Ultrapure ClO_2_ solution (3,000 ppm) was prepared by electrolysis of NaClO_2_, according to the following reaction:

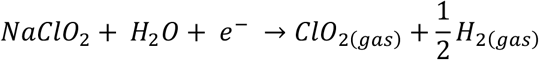

The solution was kept at 4 °C and was used within 5 days of production. Two working solutions were prepared with sterile 0.9% saline solution: a high dose (HD, 300 ppm) and a low dose (LD, 30 ppm). Final concentration was verified using test strips (LaMotte Insta-Test 3002) which allowed qualitative detection between 0 and 500 ppm.

We used 30 10-day old SPF RossxRoss chick embryos (Pilgrims, Mexico) incubated at 38°C and 65-70% humidity [48]. Five chick embryos were assigned to each of the six experimental groups at random and treated according to their group (Table 1). Total sample size in the experimental group was determined in order to achieve power 80% (type II error: 0.20) with allocation ratio 2/1, effect size of 0.7 and a significance level < 0.05. Embryos were placed at random in the incubator to avoid potential variations in temperature and humidity between the edges and the centre of the incubator. All embryos were inoculated via the chorioallanotic sac, as recommended for avian coronavirus replication [49, 50]. Details on inoculation can be seen as Supplementary Materials.

Seven days after inoculation, embryos were sacrificed according to guidelines on humane treatment and euthanasia of chick embryos over 13 days of age (AVMA 2020). The embryos were placed at 4 °C for 4 h to ensure blood coagulation and avoid contamination of the allantoic fluid [48]. The eggshell over the air chamber was disinfected with 70% ethanol and 3.5% iodine [48] and removed together with the chorioallantoic membrane. Allanotic fluid (750 μl) was collected and stored at −70 °C.

Each embryo was cleaned with distilled water to remove any remaining albumin, vitellus and amniotic fluid. Body mass was determined with a precision scale (±0.1 mg). Embryonic axis length was measured with digital callipers (± 0.1 cm). The body was examined for macroscopic lesions typically caused by avian coronavirus, such as cutaneous haemorrhages, stunting, curving, urate deposits in the kidney and feather alterations [49, 51]. Chick embryos were considered to have died during the experiment when there was evidence of disconnected or detached blood vessels of the chorioallantoic membrane or their organs exhibited imbibition of haemoglobin. Further evidence of mortality was determined by examining the presence of autolysis at microscopic examination.

Both femurs were dissected and their length was determined and averaged with a digital calliper as a surrogate measure of chick development. Samples of all organs and tissues were collected, including those known to be affected by the Massachusetts and Connecticut strains of avian coronavirus, namely trachea, lung, proventricle, duodenum, liver, kidneys and the Bursa of Fabricius [27, 49, 52, 53], and those reportedly affected by exposure to ClO_2_ [13, 54, 55].

Samples were cut longitudinally, immersed in 10% buffered formalin (pH 7.4), paraffin-embedded and sectioned at 3 μm, stained with haematoxylin-eosin and observed under the microscope at 40X and 100X. To assess haematopoietic status, bone marrow preparations were used for a 200-cell differential count to classify the marrow precursors and to determine the myeloid:erythroid (M/E) ratio for each embryo. This ratio was obtained by dividing the sum of all the granulocytic cells by the sum of all the erythrocytic cells [56].

The virus was quantified in the allantoid fluid with an RT-qPCR protocol that targeted a 100 bp fragment of the avian coronavirus N gene with primers IBV-pan_FW-1 (5′-CAG TCC CDG ATG CNT GGT A) and IBV-pan_RV (5′-CC TTW SCA GMA ACM CAC ACT)[57]. Details on the protocol can be seen as Supplementary Materials. We used a cDNA sample extracted from 500 μL of the vaccine as a positive control (the vaccine contained 10^4^ mean embryo infective dose (EID_50_)/mL of coronavirus strains Massachusetts and Connecticut). A linear regression was used to calculate the correlation coefficient (R^2^ = 0.99) and the slope value (b = −5.062) of the RNA copy number and Cq values using a 10-fold dilution of the vaccine (10^4^ to 10^−1^; see standard curve in Supplementary material). The number of viral RNA copies (hereafter viral load) was determined by comparing the Cq against this standard curve [57].

Contingency tables were built to investigate differences in the proportion of dead *vs*. live chick embryos between experimental groups. The risk of developing virus-related lesions between treated and un-treated embryos was calculated using Fisher exact tests. Body mass, morphometrics, viral load, and lesion severity were compared among groups by one-way ANOVA and post-hoc Tukey HSD tests. All analyses were performed in R version 3.6.3 [58].

## Acknowledgments

We thank Luis A. Soto-García and Karla Zamora y Cuevas for their help with photographic and video documentation of the experiment.

## Author contributions

K.A-W. and T.G-G. conceived the idea. K.A-W. supervised the experiments and wrote the manuscript. X.Z-E. performed all gross examinations and histopathology analyses. C.D-S. conducted molecular assays and artwork. M.B-M. and F.G-D performed the inoculation experiments and assisted during necropsies. K.A-W. and L.P-V conducted the statistical analyses. All authors participated in the discussion of results.

## Highlights

- Chlorine dioxide (ClO_2_) reduces mortality in chick embryos inoculated with avian infectious bronchitis coronavirus.
- Viral RNA was 2.4-fold lower in the embryos that were treated with ClO_2_.
- Microscopic lesions were less severe in ClO_2_-treated embryos.
- There was no evidence of toxicity caused by ClO_2_ at the doses used.

