## Supplementary material for "Evaluation of the antiviral effect of chlorine dioxide (ClO_2_) using a vertebrate model inoculated with avian coronavirus"

Zambrano-Estrada, X.*et al*.

### Supplementary materials and methods

Embryos were placed at random within the incubator to avoid any slight variation in humidity and temperature that could affect results. Prior to starting the experiment, all embryos were candled to ensure their viability, and were examined daily to search for evidence of death (presence of blood ring or lack of visible eggshell membrane blood vessels). Before inoculation, the eggshell was disinfected with 70% ethanol and 3.5% iodine (Guy, 2015). Using the tip of sterile scissors, a hole was drilled in the eggshell over a 1 $cm^2$ transparent tape film and a sterile 1ml syringe with a 28-g 5/16" needle was used to administer the treatment directly into the allantoic cavity as indicated in Table 1. All procedures were carried out aseptically. After inoculation, the drilled hole was sealed with a drop of glue and the embryo was returned to the incubator. Inoculation of every embryo was performed by a single person to ensure experimental variation was kept at a minimum. Embryos were candled daily to determine mortality. As there was no death recorded within the first 24 h post-inoculation, none of the embryos were discarded from the experiment.

Briefly, RNA was extracted by adding 700 µL of Trizol (Invitrogen) to 500 µL of each sample. The sample was incubated for 10 min at ambient temperature. Next, 200 µL of chloroform were added, mixed by inversion, and centrifuged at 12,000 g for 10 min at 4 °C. The aqueous phase was transferred to a sterile microtube, and 500 µL of isopropanol were added. The sample was centrifuged at 12,000 g for 10 min at 4 °C prior to decanting the supernatant. The pellet was washed with 500 µL of cold 70% ethanol, centrifuged at 7,500 g for 5 min at 4 °C and the supernatant was removed completely. The dry pellet was resuspended in 12 µL of nuclease-free water and centrifuged at 7,500 g for 1 min at 4 °C. Quantity and quality of the extracted RNA was examined in a Nanodrop (Qiagen) spectrophotometer and all samples were diluted to 35 ng/mL RNA before retrotranscription.

cDNA synthesis was performed with oligo dT and MLV reversotranscriptase (Invitrogen)according to the manufacturer's instructions. Briefly, we used 8 µL of RNA, 1 µL of Oligo DT (50 ng/µL), 1 µL of 10 mM dNTP and 2 µL of nuclease-free $H_2O$. The reactions were incubated at 65 °C for 5 min and placed on ice for 1 min. Seven µL of the kits cDNA synthesis mix, which contains 4 µL 5X First-Strand Buffer, 2 µL 0.1 M DTT and 1 µL RNAse and DNAse free $H_2O$, were added prior to mixing by pipetting and incubating at ambient temperature (~25 °C) for 2 min. We added 1 µL pf SuperScript II RT, mixed by pipetting and incubated at ambient temperature for 10 min. The reactions were incubated at 42 °C for 50 min and inactivated at 70 °C for 15 min. All cDNA samples were diluted 1:10 prior to running the qPCR assays and were stored at -20°C.

Quantitation was done with SYBR Green qPCR (Qiagen) in a real-time thermal cycler (CFX connect, Biorad) under the following protocol:  45 °C for 10 min, initial denaturation at 95 °C for 10 min and 35 cycles of 95 °C for 15 s, 52 °C for 15 s, and 68 °C for 30 s, with a final extension step at 68 °C for 10 min.

**Evaluation of the antiviral effect of chlorine dioxide ($ClO_2$) using a vertebrate model inoculated with avian coronavirus**

Zambrano-Estrada, X.*et al*.

### Supplementary figures

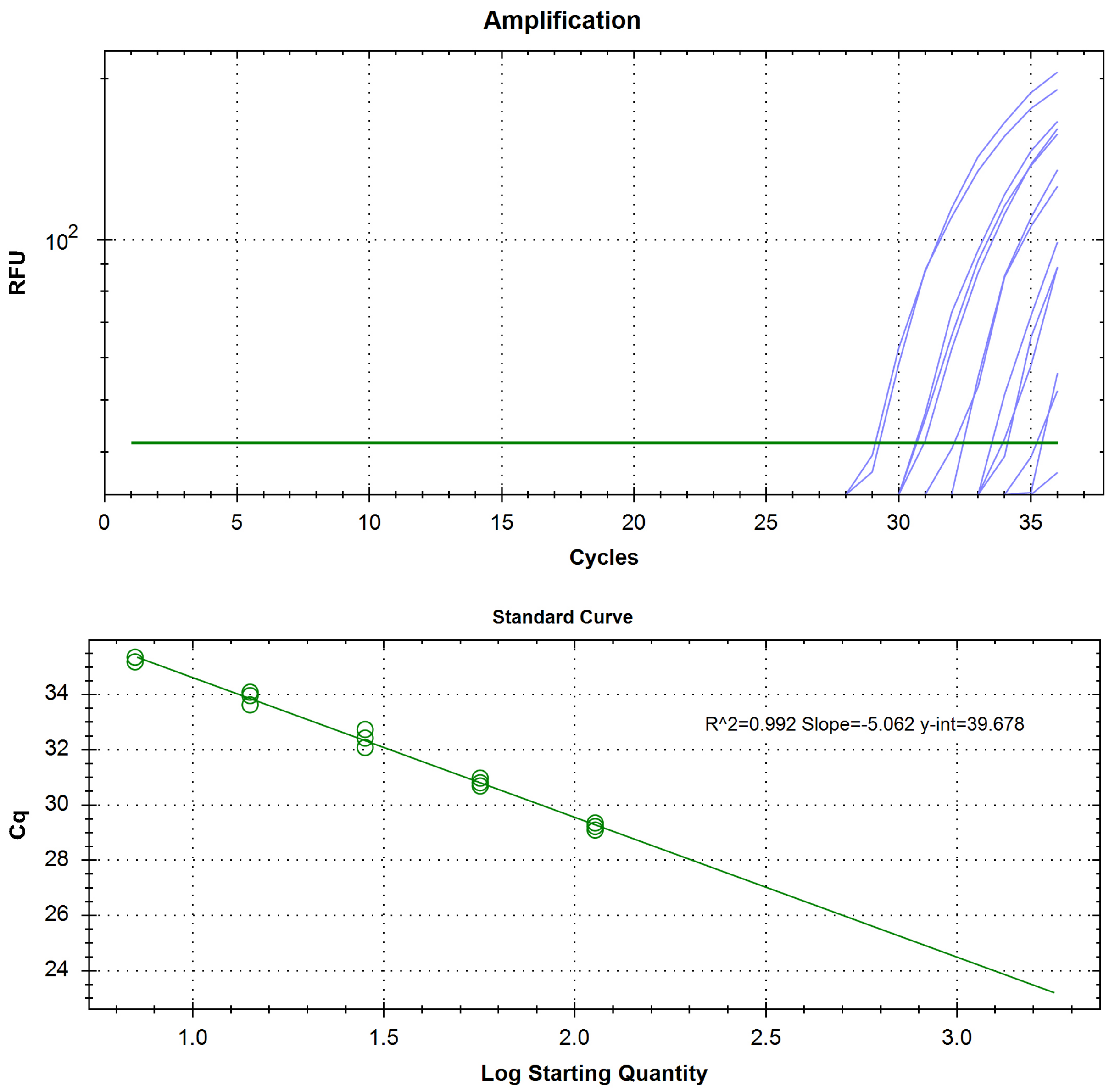


Avian infectious bronchitis coronavirus (IBV) N gene-specific RT-qPCR based on serial 10-fold dilutions of transcribed viral RNA ($10^4$ – $10^{-1}$) extracted from the live attenuated vaccine strains. The linear regression analysis of the viral RNA copy number and Cq values showed a correlation coefficient of 0.99 and slope value (b) of -5.062.

**Evaluation of the antiviral effect of chlorine dioxide ($ClO_2$) using a vertebrate model inoculated with avian coronavirus**

Zambrano-Estrada, X. *et al*.

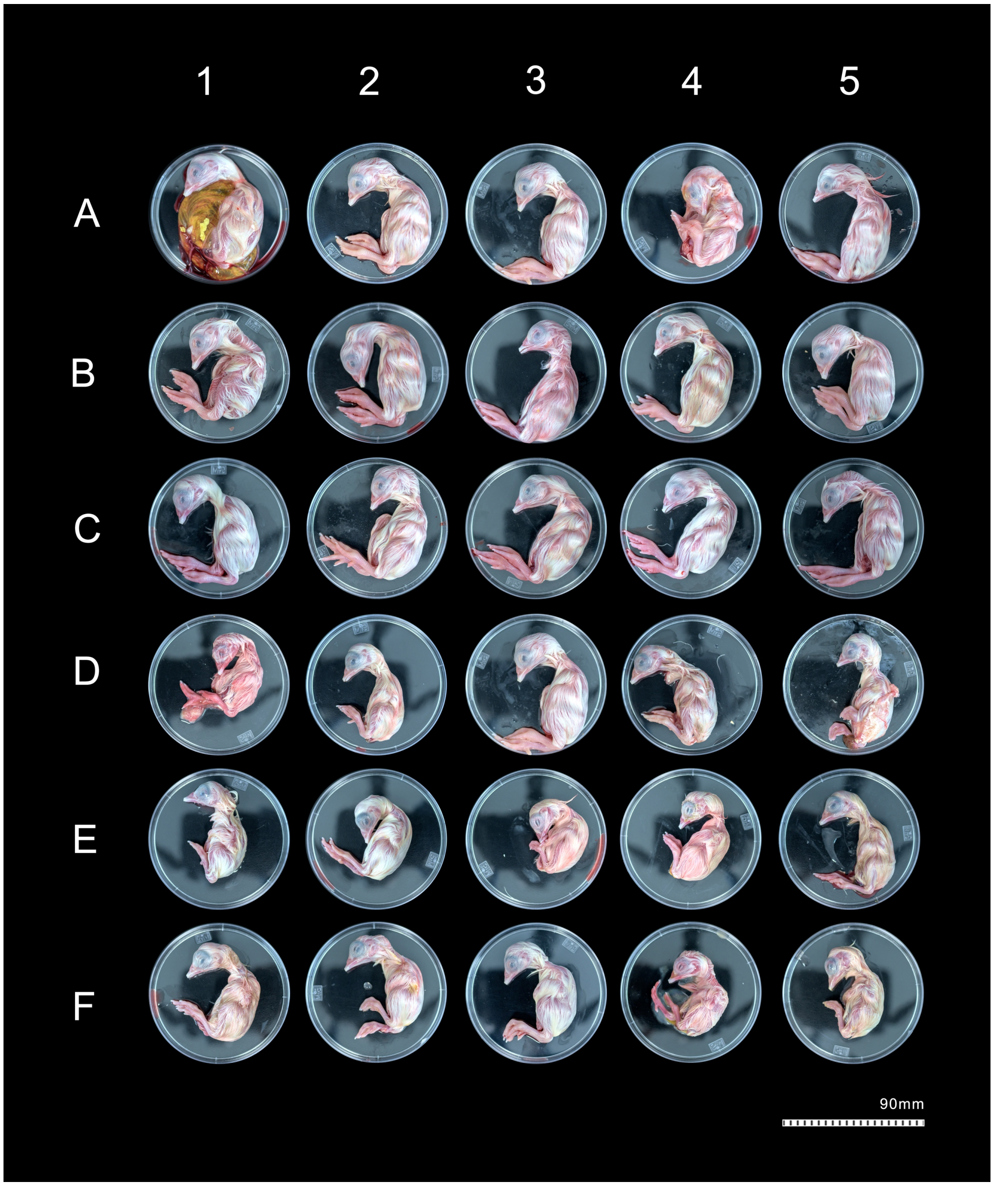


Photographs of the chick embryos at day 7 post-inoculation. A: Experimental control (embryos were administered 200 µl of sterile 0.9% chloride solution), B: Low dose of $ClO_2$ (100 µl of sterile $ClO_2$ solution (30 ppm) and 100 µl of sterile 0.9% chloride solution), C: High dose of $ClO_2$ (100 µl of sterile $ClO_2$ solution (300 ppm) and 100 µl of sterile 0.9% chloride solution), D: 100 µl of resuspended avian coronavirus vaccine and 100 µl of sterile $ClO_2$ solution (30 ppm), E: 100 µl of resuspended avian coronavirus vaccine and 100 µl of sterile $ClO_2$ solution (300 ppm), F: 100 µl of resuspended live attenuated avian coronavirus vaccine (Bron Blen® Merial, containing 104 of mean embryo infective dose ($EID_{50}$)/mL of coronavirus strains Massachusetts and Connecticut) and 100 µl of sterile 0.9% chloride solution.
